# Elucidating the role of c-di-AMP in *Mycobacterium smegmatis*: phenotypic characterization and functional analysis

**DOI:** 10.1101/2022.03.25.485789

**Authors:** Vikas Chaudhary, Aditya Pal, Mamta Singla, Anirban Ghosh

## Abstract

Cyclic-di-AMP (c-di-AMP) is a newly discovered secondary messenger molecule that plays a critical role in monitoring several important cellular processes, especially in several Gram-positive bacteria signal transduction pathways. In this study, we seek to unravel the physiological significance of the molecule c-di-AMP in *Mycobacterium smegmatis* under different conditions, using strains with altered c-di-AMP levels: c-di-AMP null mutant (*ΔdisA*) and a c-di-AMP over-expression mutant (*Δpde*). Our thorough analysis of the mutants revealed that the intracellular concentration of c-di-AMP could determine many basic phenotypes such as colony architecture, cell shape, cell size, membrane permeability etc. Additionally, it was shown to play a significant role in multiple stress adaptation pathways in the case of different DNA and membrane stresses. Our study also revealed how the biofilm phenotype of *M. smegmatis* cells are dependent on intracellular c-di-AMP concentration. Next, we checked how c-di-AMP contributes to antibiotic tolerance characteristics of *M. smegmatis*, which was followed by a detailed transcriptome profile analysis to reveal key genes and pathways regulated by c-di-AMP in mycobacteria.

## Introduction

Bacterial second messengers have been shown to modulate diverse cellular functions in different bacteria; notified examples of such messengers include (p)ppGpp, c-di-GMP, and relatively new signaling molecule c-di-AMP (1,2). The mere presence of different, yet structurally related molecules makes everybody curious to know the regulatory and functional basis of these messenger molecules under normal growth conditions and more importantly during stress adaptation. So far, c-di-AMP’s role in many Gram-positive bacteria physiology is partially known which ranges from K^+^ transport, DNA repair, cell wall metabolism, osmotic balance maintenance, antibiotic resistance, virulence, etc (3–7). In some cases, it is also shown to be involved in inducing the type I interferon response in mammalian hosts (8, 9). Decade-long research on c-di-AMP in mycobacteria has generated limited information (10–15) regarding the specific physiological functions of c-di-AMP and underlying mechanisms, which has entailed a thorough investigation of the role of c-di-AMP in *Mycobacterium smegmatis*, with and without any external stresses. In *M. smegmatis*, c-di-AMP is synthesized from condensation of two ATP molecules by enzyme DNA integrity scanning protein A (DisA) and degraded by enzyme Phosphodiesterase (Pde) into pApA. Though it was already known that the c-di-AMP level is constitutively maintained across all growth phases at a minimum concentration (10) and some basic phnotypes related to c-di-AMP in *M. smegmatis* was learntlike, further understanding regarding how the complete lack and especially the overproduction of the c-di-AMP messenger affecting stress tolerance, antibiotic sensitivity and biofilm phenotype was needed to be studied in a comprehensive and interlinked manner. We found that modulating intracellular concentrations of c-di-AMP could alter a few basic phenotypes such as cell size, cell shape, colony morphology, as well as, some surface-related properties including cellular aggregation, sliding motility, and Biofilm/Pellicle formation. Our Phenotypic Microarray (PM) data indicated differential drug susceptibility/resistance profile of the mutant strains compared to Wild-type (WT), which involves antibiotics from all major classes; these observations were subsequently confirmed by Disc inhibition and Minimum Inhibitory Concentration (MIC) assay. Further, a genome-wide transcriptome analysis (by RNA-seq) of the mutants indicated few possible cellular mechanisms behind distinctive antibiotic responses and it also highlighted critical metabolic functions and cellular pathways regulated by c-di-AMP *in vivo*.

## Materials and Methods

### Bacterial strains, media and growth conditions

A list of all the strains and plasmids used for this study is provided in Table S2 in the supplemental material. *M. smegmatis* MC^2^155 (WT) and its knockout variants *ΔdisA & Δpde* ; and their respective complemented strains (*ΔdisA+p*DisA*) & (Δpde+p*Pde) were grown in Middlebrook 7H9 broth (MB7H9; Difco) with 2% (w/v) glucose as a carbon source and 0.05% (vol/vol) Tween 80 at 37°C. The antibiotics kanamycin and hygromycin were used at a concentration of 25 μg/ml and 50 μg/ml respectively.

### *M. smegmatis* deletion mutants’ construction

To understand the role of c-di-AMP in the physiology of *M. smegmatis*, *disA* and *pde* genes were deleted by the allelic exchange as described previously (16, 17). A recombination cassette was constructed to delete both the genes individually and transformed in competent WT cells. The sucrose-resistant, gentamicin-sensitive, and kanamycin-resistant colonies were selected at 39°C for further analysis. Disruption of the gene and the recombination event were verified by PCR (showing increase in band size due to insertion of kanamycin cassette) using genomic DNA as a template, followed by sequencing.

### Complementation of Δ*disA*_Msm_ and Δ*pde*_Msm_

Functional copies of *disA* and *pde* genes were amplified from *M. smegmatis* MC2155 genomic DNA as a template using primers. The amplicon was digested with restriction enzymes (table S3) and cloned into the *E. coli-*mycobacterium shuttle vector pMV361, pre-digested with same enzymes. The final constructs pMV361_*disA* and pMV361_*pde* were electroporated in Δ*disA and Δpde* strain respectively and screened on hygromycin 7H9 agar plate supplemented with 2% glucose and 0.05% Tween.

### Construction of *disA* point mutants by Site-directed mutagenesis

The Site-directed mutagenesis (SDM) technique was adopted to generate different point mutants of *disA* and was performed by a standard procedure (18). The PCR products containing the required nucleotide substitution were transformed into *E.coli* DH5α and the resulting plasmid containing the desired missense mutation were confirmed by sequencing. The final construct pMV261-*disA*(D84A) and pMV261-*disA*(R353A, R356A) were used as templates to amplify respective *disA* mutant alleles and subcloned in mycobacterium shuttle integrative vector pMV361 enzymes using same restriction enzymes and electroporated in Δ*disA* strain and screened on hygromycin 7H9 plate supplemented with 2% glucose and 0.05% Tween. All the primers used to make deletion mutants and plasmid constructs are listed in Table S3.

### Colony architecture and morphology

As described earlier (19), 20 μl of the early stationary phase grown culture of *M. smegmatis* was spotted in the middle of the MB7H9 agar plates supplemented with 2% glucose and incubated at 37°C for 8 to 12 days. Post incubation, images of the individual colonies were taken. For colony morphology, mid-log phase cultures were diluted 1:1000 and spotted four microliters on MB7H9 agar plates supplemented with 2% glucose and incubated at 37°C for 3 days. Post incubation, images of the individual colonies were taken.

### Transmission electron microscopy

Transmission electron microscopy was carried out by adapting the protocol from Ghosh et al (20). Mid-log-phase *M. smegmatis* cultures were washed with PBST (Phosphate-buffered saline tween) twice and then 3 μl of the washed culture was drop cast on carbon-coated copper grids (Tedpella) and let stand for 10 minutes at room temperature. The sample containing grid was blot dried using the Whatman filter paper and was negatively stained using 0.5% of uranyl acetate solution, then allowed to air dry. Then the stained grids were imaged with Talos L120C transmission electron microscope (ThermoFisher) operated with 120 kV at room temperature.

### Scanning electron microscopy

The procedure of scanning electron microscopy was adapted from a previously published study (21). Mid-log-phase *M. smegmatis* cultures were fixed with 2% glutaraldehyde for 30 minutes at room temperature and then stored at 4°C for 4 hours. After that, cells were washed with Phosphate buffer (0.1M, pH-7.4) 3 times and resuspended in the same buffer. Samples were dehydrated through ethanol series 30%, 50%, 70%, 90% and 100 % at 4°C. A thin film of the sample was applied on a coverslip, air-dried for 20 minutes followed by sputter coating with gold particles; and observed under Zeiss Merlin VP FESEM with SE2 detector.

### Biofilm formation assay

As described before (22), biofilms were grown in the Sauton’s fluid base medium supplemented with 2% glucose as a carbon source. 1.5 ml. of the media was poured into each well of a 24-well plate and 1.5 μl of the washed (twice) stationary phase cells were added into each well and the plate was incubated at 37°C in a humidified incubator for 4 days and the images were recorded under white light in a set of 3 biological replicates.

### Sliding motility assay

Sliding motility assay was adapted from Martinez et al (23). Briefly, the cells were grown till OD_600_ 1.2 then diluted to OD_600_ 0.4 in PBST buffer. 3 μl of the diluted cultures were spotted in the middle of MB7H9 plates solidified with 0.3% agarose without any carbon source. The plates were incubated at 37°C for 2-3 days in a humidified incubator. Motility was measured by the diameter of the halo growth.

### Cellular aggregation assay

Cellular aggregation assay was adapted from Deshayes et al (24). Briefly, early stationary phase cells were centrifuged at 10,000g for 1 minute. CFU spotting was done with both supernatant and pellet. The aggregation percentage was calculated by using the CFU values of supernatant and pellet.

### Ethidium bromide influx Assay

Influx assay was adapted from Zhang et al. .Briefly, *M. smegmatis* cultures were grown at 37°C in MB7H9 medium to an OD_600_ of 0.9-1. Cells were washed twice with phosphate buffer saline (PBS) with 0.05% tween80, then resuspended in 1/3^rd^ volume of the same buffer and kept at 37°C shaking for one hour to induce starvation. After that, carbonyl cyanide m-chlorophenylhydrazone (100 μM) was added (acts as efflux pump inhibitor) and cells were further incubated for 30 minutes in the same condition. Next, Ethidium bromide (0.5 μg/ml f.c.) was added and cells were immediately taken into 96 well black plates. To quantify the accumulation of EtBr, fluorescence was checked for the next one hour (with a 5-minute interval between two readings) in Varioskan Flash multimode reader (Thermo Fisher Scientific) at 530 nm and 590 nm wavelength for excitation and emission respectively.

### Pellicle formation assay

As described previously (26), stationary-phase cultures were diluted to an OD_600_ of 0.03 in LB media and kept standing for at 37°C without any disturbances and then imaged after 5 days.

### Stress tolerance assays

*M. smegmatis* strains were grown until the mid-log phase and then normalized to OD_600_ 0.7. The bacteria were diluted by 10-fold serial dilutions and 4 μl of cells from each dilution were spotted into MB7H9 agar medium. After the drying of the spots, the plate was exposed to UV irradiation (0.15 mJ/ cm^2^) (27) and then incubated at 37°C for 3 days in the dark. For SDS treatment, OD_600_ 0.7 cells were exposed to 0.05% SDS (f.c.) for 4 hours and then spotted on MB 7H10 agar plates. During MMS treatment, mid-log phase cultures were centrifuged and resuspended in an equal volume of PBST and treated with 0.5% MMS (f.c.) for 20 minutes at 24°C then spotted (10-fold serial dilution) on MB 7H10 agar plates (11). Survival percentage was calculated as the percentage of CFU remaining after exposure to the respective stresses compared to the CFU in an untreated condition.

### Phenotypic microarray analysis

Phenotypic microarray (PM) assay was performed using the protocol of Gupta *et al* (22). In brief, the transmittance of the logarithmic phase bacterial cultures was adjusted to 81% and tetrazolium violet was added to a final concentration of 0.01%. 100 μl of the cultures were inoculated into the wells of PM plates 1 to 20 coated with chemicals/antibiotics and kept in the Omnilog incubator (Biolog) at 37°C for 4 days. The dye reduction values were converted to the area under the curve (AUC) by using the parametric software module of Biolog Omnilog software. The AUCs for the strains were compared and overlaid as the test strain versus reference strain AUC.

### MIC (Resazurin Microtiter Assay Plate, REMA) assay

MIC values were estimated using REMA adapted from an earlier protocol (28). In brief, the transmittance of the culture was adjusted to a McFarland turbidity standard of 1 and then diluted to 1:10. Next, 196 μl portions of the diluted culture were inoculated into 96 well microtiter plates containing a 2-fold serial dilution of the antibiotics (4 μl). Plates were sealed and incubated for 37°C. After 36 hours, 30 μl of 0.01% resazurin dye was added to each well, and plates were further incubated for 4-6 hours. The color of the resazurin changed from blue to pink due to bacterial growth. The MIC was determined as the minimum antibiotic concentration at which the resazurin dye did not change the color.

### Disc diffusion assay

As previously described (29), *M. smegmatis* strains were grown until the mid-log phase and 100 μl of the OD_600_ 0.7 cultures were spread on MB 7H10 agar plates and left for drying. After that, discs soaked with 10 μl of erythromycin (50 mg/ml) and ciprofloxacin (2.5 mg/ml) stocks were aseptically kept in the middle of the plate using a sterile tweezer. Plates were further incubated at 37°C for 3 days and the zone of inhibition (cm^2^) was calculated as a measurement of antibiotic sensitivity.

### Transcriptome analysis by RNA-Seq

As described earlier (30), *M. smegmatis* cells were grown till early exponential phase (OD_600_~0.6), washed twice with PBS and dissolved in RNA later solution with TRIzol (Invitrogen) and placed on ice. Subsequent steps of RNA extraction, DNA synthesis, library preparation, and alignment were carried out at Clevergene, Bangalore, India. Briefly, the cells were mechanically disrupted by bead beating and the supernatant was collected for RNA extraction using HiMedia HiPurA extraction kit and finally eluted with 15ul of RNase free water. The RNA quality assessment was done using RNA HS ScreenTape System (Catalog: 5067-5579, Agilent) and the RNA concentration was determined on Qubit® 3.0 Fluorometer (Catalog: Q33216, ThermoFisher Scientific) using the Qubit™ RNA HS Assay Kit (Catalog: Q32855, ThermoFisher Scientific). Next, 500ng of total RNA was taken for rRNA depletion using QIAseq FastSelect −5S/16S/23S Kit (Catalog: 335925, Invitrogen) according to the manufacturer’s protocol. NEBNext Ultra II RNA Library Prep Kit (Catalog: Catalog: E7775S, New England Biolabs) for Illumina was used for the library preparation. The enriched transcriptome was chemically fragmented in a magnesium-based buffer at 94°C for 10 minutes. The fragmented samples were primed with random hexamers, and reverse transcribed to form cDNA and the first-strand cDNA reactions were converted to dsDNA. Furthermore, the DNA was amplified by 12 cycles of PCR with the addition of NEBNext Ultra II Q5 master mix, and “NEBNext® Multiplex Oligos for Illumina” to facilitate multiplexing while sequencing. After the library quantification and validation, the sequence data were generated using Illumina HiSeq. After removing adapter sequences and low-quality bases, the processed reads were mapped on the reference genome *Mycolicibacterium smegmatis* MC^2^155 for further analysis of differential expression, gene ontology and pathway enrichment.

## Results

### Modulating intracellular c-di-AMP concentration affects basic phenotypes

To directly study the role of c-di-AMP in *M.smegmatis* basic physiology, we used two deletion mutants*: M. smegmatis ΔdisA* and *M. smegmatis Δpde*, which corresponds to c-di-AMP null mutant and over-expressing mutant strains respectively. As a part of strain validation, apart from PCR **(Fig.S1)** and sequencing confirmations assuring respective gene deletions, we measured the c-di-AMP concentration in both the deletion mutants along with WT *M. smegmatis* by HPLC **(Fig.S2)**. HPLC analysis indicated the lack and abundance of the c-di-AMP molecule in respective mutant strains. Once the strains were verified, we constructed individual complementation strains by cloning *disA* and *pde* genes in a single copy integrating vector pMV361 and transformed them to the corresponding knockout strains resulting in the construction of complementation strains. With these 5 strains, we first checked how the basic phenotypes which have been affected by unbalancing the steady-state homeostasis of c-di-AMP messenger, which is hypothesized to be the key factor during normal growth and stress tolerance.

First, we began our study by checking a few basic cellular phenotypes (without any stress) which were partly known (10,12) and our primary observation inferred minute growth deficiency in *M. smegmatis Δpde* strains in broth cultures and agar plates. Precisely, *M. smegmatis Δpde* strain had a minor growth defect in the exponential phase in MB7H9 media **(Fig.S3)** and formed smaller colonies with different morphology on agar plates **(Fig.1A)**. These observations were in line with previous studies where plasmid based ectopic overexpression of *disA* (pMV261-*disA*) was carried out in *M. smegmatis* (10, 38). Consequently, when we tested *M. smegmatis Δpde*+ pMV361-*pde*, we could not observe any growth defects highlighting the direct role of high intracellular concentration of c-di-AMP, and not immoderate DisA-DNA interaction behind the phenotype **(Fig.S3)**. However, *M. smegmatis ΔdisA* strain did not show any growth deficiency **(Fig.S3)** compared to WT further implying the significance of maintaining a low concentration of c-di-AMP inside cells. Similar observations with smaller colony size in a high c-di-AMP producing strain have been reported previously with overexpression constructs (10).

**Figure 1.**
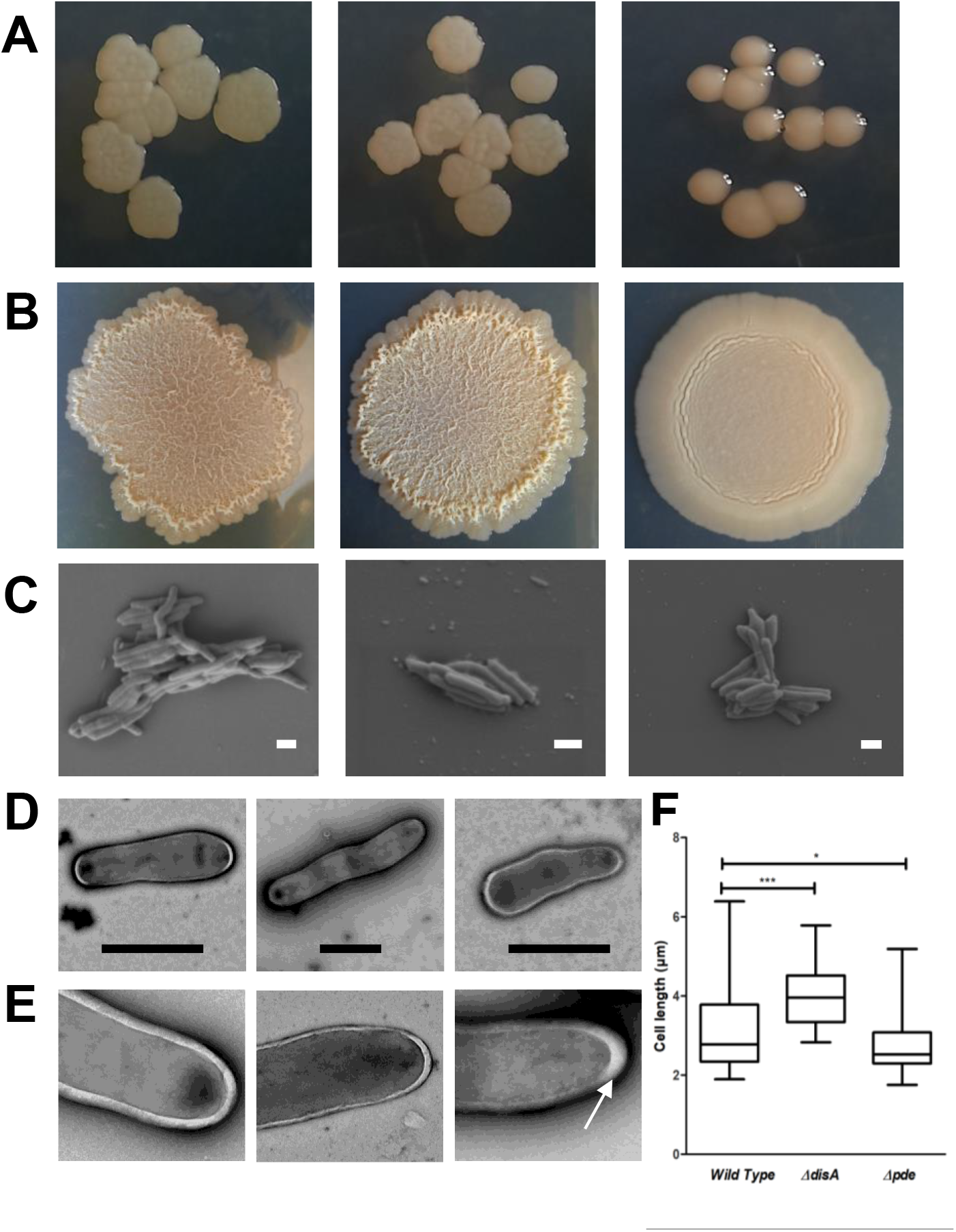
Modulating intracellular c-di-AMP concentration affects the colony characteristics of *M.smegmatis* WT(left panel), *M.smegmatis ΔdisA* (middle panel) *& M.smegmatis Δpde* (right panel): **(A)** Colony size and morphology **(B)** Colony architecture. Individual cell shape and size determination by electron Microscopy. Representative microscopic images of *M.smegmatis* WT(left panel), *M.smegmatis ΔdisA* (middle panels) *& M.smegmatis Δpde* (right panels), with SEM **(C)** and TEM **(D)** microscopy (scale bars = 2 μm) revealed the difference in cell morphology linked to intracellular c-di-AMP concentration. **(E)** Closer inspection of some TEM images revealed occasional increased outer layer thickness at poles in *M.smegmatis Δpde* strains, highlighted by white arrow; **(F)** Cell length distribution analysis of individual strains. Lengths of at least 50 cells of each strain from different electron micrographs were measured and plotted using the box plot analysis function from GraphPad Prism 5. The whiskers in the plot represent minimum and maximum values. The mean lengths were found to be 3.11 μm for the WT, 3.98 μm for the *M.smegmatis ΔdisA* and 2.70 μm for the *M.smegmatis Δpde* strain. The data was plotted using GraphPad Prism 5. *** = *P* < 0.001; ** = *P* < 0.01; *\* = P* < 0.05.

Next, we found that the colony architecture was drastically changed in the c-di-AMP overexpressing strain *M. smegmatis Δpde* compared to WT, and the phenotype was reversed in the respective complementation strain **(Fig.S4)**. The colony surface became smooth, uniform, and glossy in the *M. smegmatis Δpde* strain **(Fig.1B)**, which is possibly linked with major changes in surface properties. On the other hand, all the other strains formed characteristic *M. smegmatis* rough and dry colonies **(Fig.1B, Fig.S4)**.

Once we were certain that some of the basic phenotypes were consequentially altered, we were interested in changes in individual cell shape and size of WT and mutants. First, we did Scanning Electron Microscopy (SEM) to reveal differences in 3D cell morphology and surface properties and found that cell length was marginally increased in the case of *M. smegmatis ΔdisA* strain compared to WT, whereas in a high c-di-AMP strain (*Δpde)* cells have become shorter and thicker **(Fig.1C)**. Previously it has been reported also that altering the concentration of another second messenger such as (p)ppGpp could have an impact on *M. smegmatis* cell elongation (31). Further analysis with Transmission Electron Microscopy (TEM) suggested that *M. smegmatis ΔdisA* cells possess almost uniform cell-width and slightly reduced outer layer thickness compared to WT cells. On the other hand, *M. smegmatis Δpde* cellular width is variable and often shown to have a thicker outer layer, especially in the poles **(Fig.1E)**. Finally, we measured the individual cell lengths of different strains from the TEM image dataset, and the comparison revealed significant cell length variation across strains as a function of c-di-AMP concentration **(Fig.1F)**. Though significant differences in terms of cell length, shape, and outer layer thickness were observed linked to varying c-di-AMP concentrations, the exact pathway/ mechanism needs to be studied in the future.

### Surface properties are significantly altered in high c-di-AMP strain

Visible differences in colony architecture of *M. smegmatis Δpde* strain demanded a further investigation on other cell surface properties. Hence, we quantified the level of cellular aggregation by a previously described protocol (22) and found that the intracellular c-di-AMP concentration was directly correlating with the aggregation level of cells, where *M. smegmatis Δpde* tends to form more cellular aggregates and *M. smegmatis ΔdisA* strain formed lesser aggregate compared to WT. As expected, both *disA* and *pde* complementation strain behaved like a WT **(Fig.2A)**.

**Figure 2.**
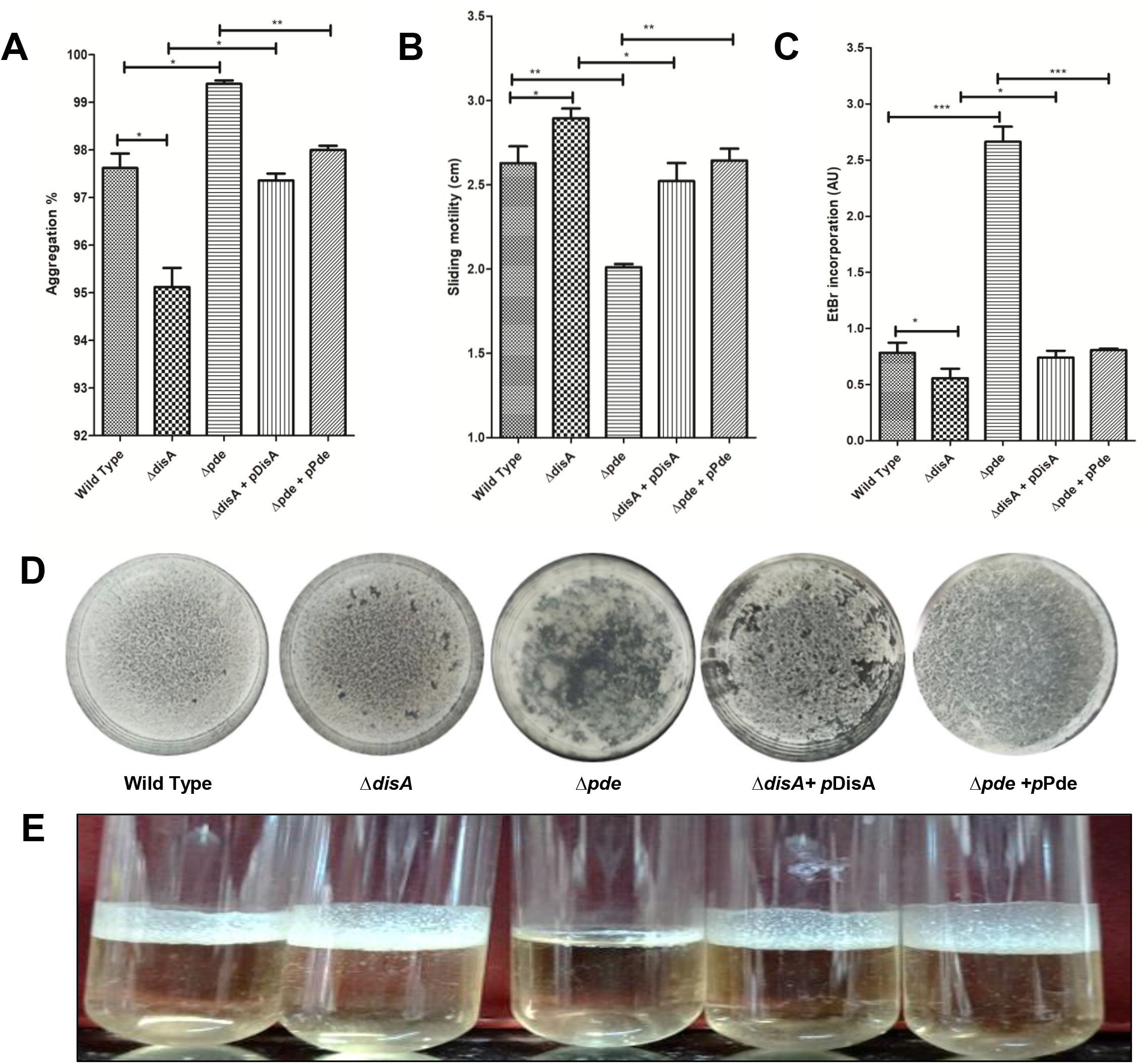
Estimation of cell surface-associated Phenotypes by comparison of **(A)** Cellular aggregation profile(%), **(B)** Sliding motility (cm.) and **(C)** EtBr incorporation (AU) in *M.smegmatis* WT, deletion mutants and the respective complementation strains. Error bars indicate the standard deviations from three independent experiments. **(D)** Biofilm formation phenotypes showed *M. smegmatis* WT and *M.smegmatis ΔdisA* forms a thick biofilm in 24 well plates at 37 degrees after 4 days, whereas *M.smegmatis Δpde* strain is defective in biofilm formation panels but the phenotype is restored in the complementation strain. **(E)** Pellicle formation was estimated at the air-liquid interface of the standing LB culture of different strains and *M.smegmatis Δpde* strain was found to be significantly compromised in forming a pellicle.The graphs were plotted using GraphPad Prism 5. *** = *P* < 0.001; ** = *P* < 0.01; *\* = P* < 0.05.

Next, we checked the surface hydrophobicity as a function of glycopeptidolipids (GPL) of individual strains in terms of sliding motility on agarose plates using a previously described method. Here we found that *M. smegmatis ΔdisA* strain was visibly more motile than WT, on the other hand, *M. smegmatis Δpde* strain had significantly lost motility compared to WT **(Fig.2B)**. A major change in cellular aggregation and sliding motility was also reported before using *M. smegmatis* + pMV261-*disA* strain, but that could be a indirect effect of a severe growth deficiency of the strain reported in the same study (38). Moreover, just by overexpressing DisA (a dual function protein), it was not clear if the observed phenotype was due to high c-di-AMP concentration or increased DisA-DNA interaction. In our study, we showed *pde* complementation strain’s aggregation profile and sliding motility pattern remain unchanged like WT (**Fig.S5)**, which further pointing towards a direct link between intracellular c-di-AMP concentration with aggregation and motile property of *M. smegmatis*. Often this change in surface motility has been linked to different glycopeptidolipids (GPL) fractions in the cell wall (32, 33) and from this result, it is evident c-di-AMP could possibly play a role in cell wall metabolism. As it has been previously mentioned that c-di-AMP responsive transcriptional factor DarR is responsible for regulation of fatty acid biosynthesis in *M. smegmatis* (12), the drastic change in surface properties could be due to the alteration of fattyacids composition in cellwall and membrane.

Our hypothesis regarding altered surface properties linked with envelope structure and porosity was explored by the Ethidium bromide (EtBr) influx assay. It was found that EtBr influx was almost 3 fold higher in *M. smegmatis Δpde* strain from WT, whereas no difference was observed with *M. smegmatis ΔdisA* strain **(Fig.2C)**. Since this assay was done in presence of a well-known efflux pump inhibitor CCCP (34) at a non-toxic concentration, the differential uptake of EtBr highlighted the difference in membrane permeability or cell wall rigidity (5, 35) or functional regulation of particular membrane transporters by c-di-AMP, which needs to be investigated in the future.

### High c-di-AMP concentration inhibits biofilm formation in *M. smegmatis*

One of the well-studied phenotypes linked with second messengers such as (p)ppGpp and c-di-GMP is biofilm formation. We also wanted to study the possible link between c-di-AMP concentration and biofilm formation characteristics in *M. smegmatis*. Our data suggested that, though *M. smegmatis ΔdisA* strain had no visible difference in biofilm formation in 24 well plates with WT, *M. smegmatis Δpde* possess inadequate ability to form and maintain biofilm in identical conditions **(Fig.2D)**. In mycobacteria, air-liquid interphase pellicle formation is also often considered as a reliable marker for biofilm formation and we found that *M. smegmatis Δpde* strain was significantly compromised in forming pellicle when grown in glass tubes, whereas *M. smegmatis ΔdisA* strain formed slightly more pellicle than WT**(Fig.2E)**. In both the direct and indirect assays to quantify biofilm formation, the high c-di-AMP strain showed a clear difference in biofilm formation further highlighting c-di-AMP’s contribution in maintaining such complex phenotypes. The fact that there was hardly any difference between WT and *ΔdisA* strain, it clear that c-di-AMP’s presence and absence does not control biofilm properties in *M. smegmatis*, rather the intracellular c-di-AMP concentration plays a critical role in rigid biofilm formation in *M. smegmatis* whereas a closely related second messenger c-di-GMP has proved to have a negligible impact (36). Thus, any ectopic increase in c-di-AMP level severely affects the biofilm formation phenotype of *M. smegmatis* and possibly makes the cells more prone to external stresses, such as antibiotics. Hence from a bacterial cell’s point of view, it is pivotal to keep the c-di-AMP level under check.

### c-di-AMP is involved in several stress tolerance pathways

Since c-di-AMP and DisA have been previously linked with DNA damage repair in different organisms including *M. smegmatis* (37–39), we first estimated the survival of the deletion mutant strains against several genotoxic stresses.

Our data reconfirmed the previous report (11) that *M. smegmatis ΔdisA* strain is very sensitive (~30 fold) to 0.5% methyl methanesulfonate(MMS) treatment, which is known to methylate DNA bases and cause replication fork arrest. On the contrary, high c-di-AMP mutant *M. smegmatis Δpde* showed a much smaller difference with WT **(Fig.3A)**, which could indicate that the DisA scanning enzyme, rather than c-di-AMP molecule, is important to prevent MMS driven toxicity by an unexplored mechanism.

**Figure 3.**
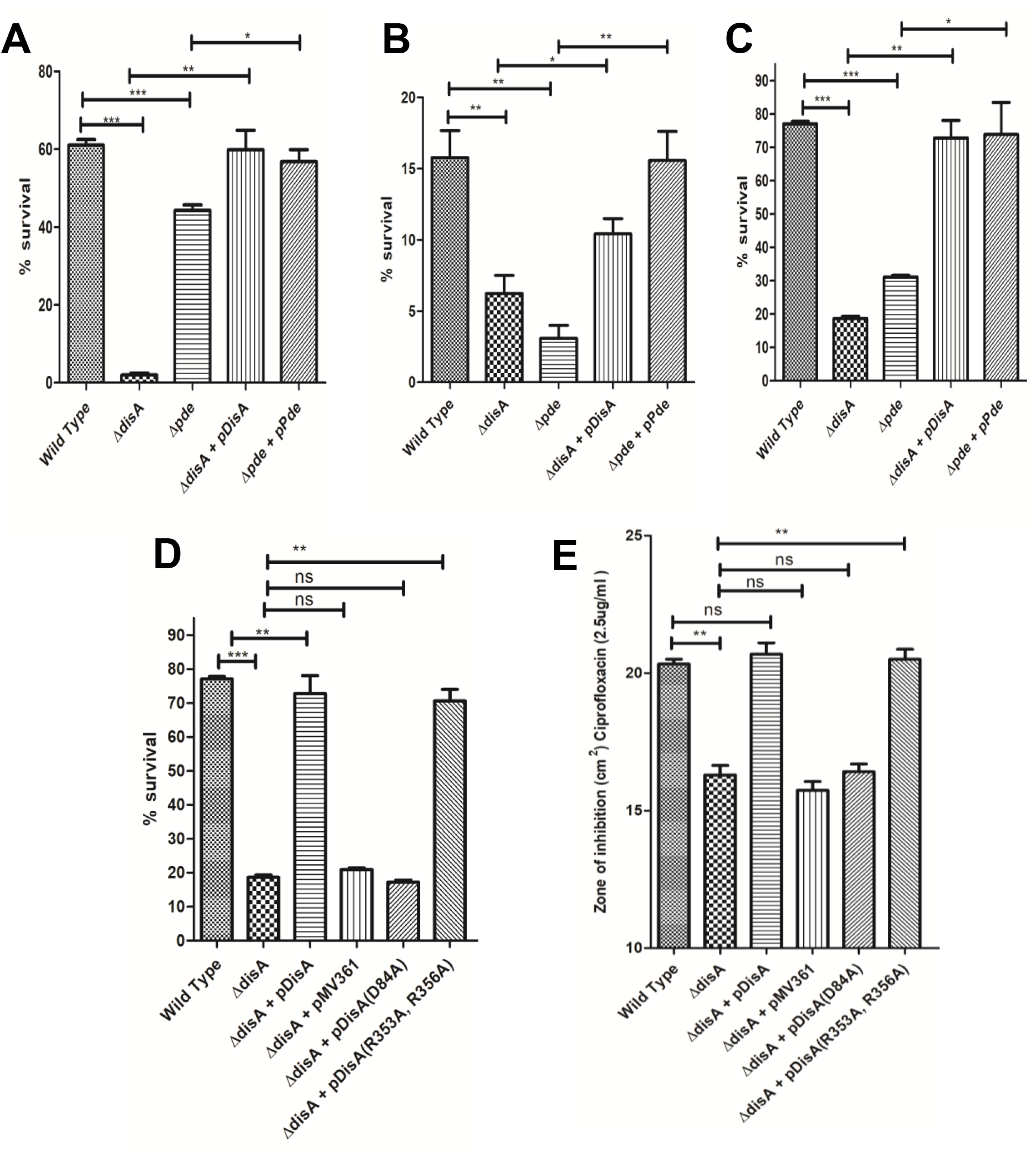
Stress tolerance assays. *M.smegmatis ΔdisA* & *M.smegmatis Δpde* strains display sensitivity to several stresses. Percentage survival was calculated after 0.5% MMS treatment **(A)**, 0.15 mJ/ cm^2^ UV irradiation **(B)**and 0.05% SDS treatment **(C)**. DisA protein’s multi-domain nature and interdomain functional independency were illustrated by SDS treatment **(D)**and disc assay predicting ciprofloxacin sensitivity **(E)**. The graphs were plotted using GraphPad Prism 5. *** = *P* < 0.001; ** = *P* < 0.01; *\* = P* < 0.05.

Next, we checked how the deletion mutants behaved under UV exposure (0.15 mJ/ cm^2^). We found that the *M. smegmatis ΔdisA* strain is ~3-4 fold more sensitive than WT. To our surprise, we found that the *M. smegmatis Δpde* strain was found to be extremely sensitive to UV stress (~5-6 fold) compared to WT **(Fig.3B)**, which is possibly linked to impeded *recA* function (11). The fact that both null- and overexpressing c-di-AMP mutants were sensitive to UV stress, apparently implied that both higher and lower concentration of c-di-AMP is detrimental for cells under such conditions.

Other than, DNA stresses we treated the mutants with ionic detergent Sodium dodecyl sulfate (SDS) and found that both *M. smegmatis ΔdisA* and *M. smegmatis Δpde* strains were found to be sensitive (20-30% survival) to 0.05% SDS compared to WT **(Fig.3C)**. Respective complementation strains showed a similar level of sensitivity (70-80% survival) with WT further suggesting how the varying concentration of c-di-AMP could make cells more prone to cell lysis possibly due to the membrane and cell wall protein denaturation **(Fig.3C)**.

Finally, as c-di-AMP has been previously studied (4, 40) to contribute to ion homeostasis and envelope stress in several Gram-positive bacteria, we checked the c-di-AMP mutants strains’ (*M. smegmatis ΔdisA* and *M. smegmatis Δpde)* response against osmotic stress (500 mM NaCl and KCl) and found that there was no visible difference observed in the tolerance/survival levels when compared to *M. smegmatis* WT and *ΔdisA* mutant, but *Δpde* mutant showed a low level of sensitivity (data not shown).

Representative spot plate pictures of each assay along with control strains were shown in **Fig.S6**.

### Identifying DisA protein’s multi-domain nature and mutational uncoupling of two activities

DisA being a multi-domain and dual-functional protein (38), we wanted to understand the specific role of different domains during stress tolerance by individually putting missense amino acid substitutions of the key residues either known or identified through bioinformatic modeling. As we found out, *M. smegmatis ΔdisA* strain has shown clear sensitivity against SDS treatment and significantly resistant phenotype to ciprofloxacin compared to WT (described by Phenotypic Microarray, MIC and disc inhibition assay later) we complemented *M. smegmatis ΔdisA* strain with different *disA* mutants (c-di-AMP synthesizing catalytic mutant and DNA binding mutant) in pMV361 integrative vector. In all our assays, *M. smegmatis ΔdisA+p*DisA(WT copy) and *M. smegmatis ΔdisA+p*Empty served as positive and negative controls respectively to check reversal of phenotype for *M. smegmatis ΔdisA* strain to WT upon complementation with different DisA point mutants (D84A:c-di-AMP synthesizing catalytic mutant (38); R353A, R356A (putative DNA binding mutant). Upon SDS treatment it was found that *M. smegmatis ΔdisA+p*DisA(D84A) behaved similarly like *M. smegmatis ΔdisA+p*Empty and failed to reverse the SDS sensitivity, whereas *p*DisA(R353A, R356A) complementation construct was able to make the *M. smegmatis ΔdisA* strain less sensitive to 0.05% SDS like *M. smegmatis ΔdisA+p*DisA(WT copy) **(Fig.3D)**. Then, we checked whether the ciprofloxacin resistance phenotype was reversed with complementation of different DisA point mutants and it was found that, *M. smegmatis ΔdisA+p*DisA(D84A) retained the resistance phenotype like *M. smegmatis ΔdisA* strain and *M. smegmatis ΔdisA+ p*DisA(R353A, R356A) strain became sensitive to ciprofloxacin like WT and *M. smegmatis ΔdisA+p*DisA(WT copy) complementation strain **(Fig.3E)**.

Both these observations led us to ascertain the fact that SDS sensitivity and ciprofloxacin resistance phenotypes of *M. smegmatis ΔdisA* strain are due to lack of c-di-AMP and not due to loss of DisA enzyme’s DNA scanning property. This mutational uncoupling study of the DisA protein by targeting single amino acid residues in either domain of the enzyme further corroborated the multi-domain characteristics and inter-domain functional independency of this important dual-function protein in *M. smegmatis*.

### Phenotypic Microarray analysis revealed a direct link between c-di-AMP concentration and differential drug susceptibility

To gain a deeper understanding of how c-di-AMP concentration plays a critical role in drug tolerance, we performed Phenotypic Microarray (PM) analysis which gave a vast platform to check antibiotic sensitivity/resistance phenomenon across various concentrations of 240 different antibiotics/chemicals in a high throughput manner (41). The growth of the respective strain in presence of a particular compound was calculated based on dye reduction values plotted as a function of time up to 96 hours. Whether the c-di-AMP null- and overexpressing-mutants were growing better or worse than WT was deduced by PM Omnilog software which could simultaneously compare the AUC (Area Under Curve) values of two strains for a specific antibiotic concentration **(Fig.S7)**. By analyzing the significant differences in AUC values between *M. smegmatis* WT and *Δpde* strain first, we shortlisted a few antibiotics (with different Mechanism of action) for which *Δpde* strain showed hypersensitivity, as evidenced by lower AUC values. Next, we compared the AUC values of the same drug concentrations (specific well in the PM plate) between *M. smegmatis* WT and *ΔdisA* strain to know if there was any similar or more importantly reverse trend was observed. Based on these considerations, our phenotypic microarray data showed that the *M. smegmatis Δpde* strain was significantly more sensitive to an array of antibiotics like Ciprofloxacin, Rifampicin, Erythromycin, and Vancomycin **(Fig.4A)**. Though the level of sensitivity differed between different antibiotics, this preliminary observation has thrown a light on how c-di-AMP concentration could imitate the multi-drug tolerance phenotype of a strain, which was not reported anywhere as per our knowledge. In the case of all 4 antibiotics, *M. smegmatis ΔdisA* strain showed moderate/ significantly higher AUC values compared to WT further corroborating essentiality for an optimum intracellular concentration of c-di-AMP. Next, we confirmed the PM observation of increased sensitivity of *Δpde* strain in two ways: Minimum Inhibitory Concentration (MIC) assay and antibiotic disc diffusion assay, to assess the effect in both liquid and solid media. To be sure, the observed drug sensitivity was indeed due to the deletion of *pde* gene, and not due to any secondary polar effect arising from the deletion of the gene we used complementation strain (*M. smegmatis Δpde +p*Pde) in our MIC and disc diffusion assays. Both the complementation strain: *Δpde+p*Pde and *ΔdisA+p*DisA showed similar tolerance levels (MIC value and zone of inhibition) for all 4 antibiotics highlighting the direct relationship between varied c-di-AMP concentration and differential drug susceptibility, whereas *M. smegmatis Δpde* strain 2-8 fold more sensitive to all 4 antibiotics compared to WT strain **(Fig.4B)**. Similarly, *M. smegmatis Δpde* strain produced a significantly bigger zone of inhibition in presence of 2.5 μg/ml. ciprofloxacin **(Fig.4C)** and 50 μg/ml. erythromycin **(Fig.4D)** discs and with the single copy complementation of the *pde* gene the sensitivity was reverted to the WT level. Though there is a common trend that high intracellular c-di-AMP concentration makes the cell more vulnerable against a different class of antibiotics, the specific molecular mechanism related to the individual mode of action of the drug remains to be investigated in the future. **Table S1**contains the list of the top 40 antibiotics/chemicals in which the area under curve (AUC) difference between WT and mutants was shown to be the highest on either side (loss and gain).

**Figure 4.**
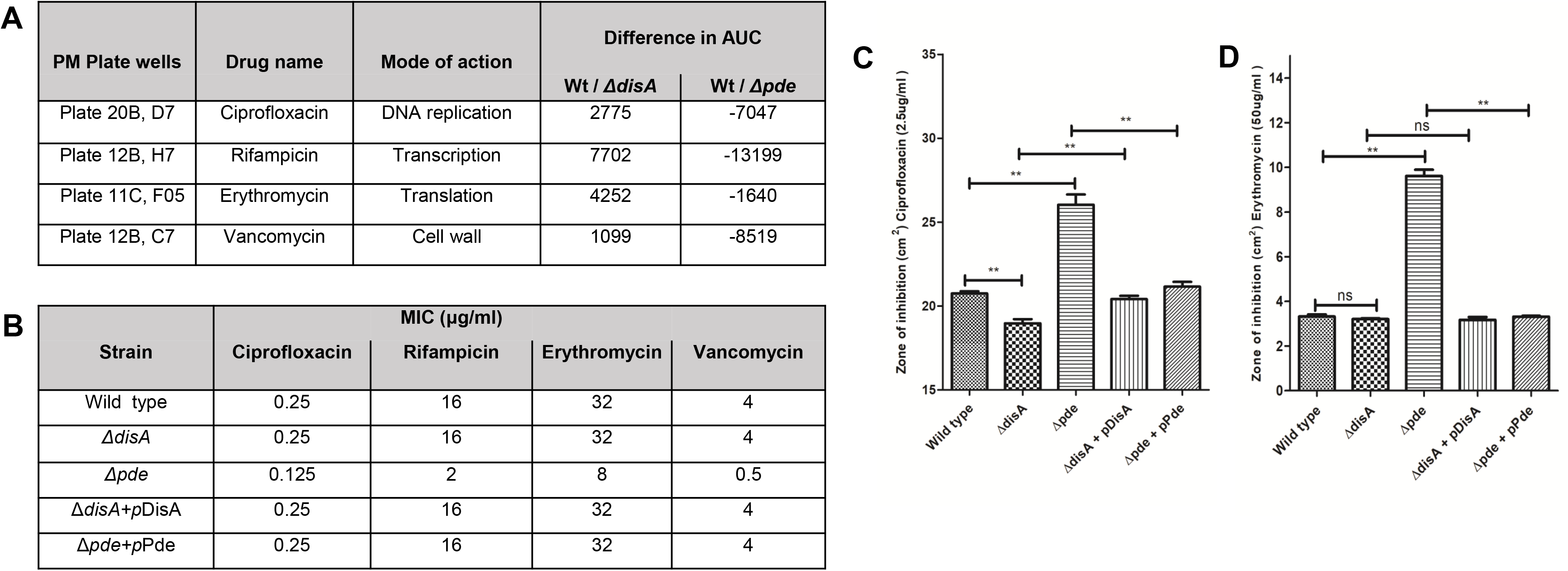
Phenotype microarray analysis in *M.smegmatis* WT, *M.smegmatis ΔdisA & M. smegmatis Δpde* strains. The strains were grown in 96-well plates in the presence of different antibiotics with tetrazolium violet as an indicator for growth. The change in the color of the dye due to respiration was measured, and the absorbance was plotted as a function of time. **(A)** Comparison of growth profiles in terms of area under curve (AUC) values. **(B)** MIC values with representative antibiotics for the Wild-type *M.smegmatis* and its isogenic variants were determined using the REMA method. Disc inhibition assay n terms of zone of inhibition were calculated with ciprofloxacin **(C)** and erythromycin **(D)**, revealed a high level of sensitivity *M.smegmatis Δpde* strain, further confirming PM observation and elucidating the direct relationship between c-di-AMP concentration and differential drug susceptibility. *** = *P* < 0.001; ** = *P* < 0.01; *\* = P* < 0.05.

### RNA-seq based transcriptome analysis highlighted key pathways regulated by c-di-AMP

After we confirmed c-di-AMP’s direct role in both stress adaptation and drug sensitivity, further insights into the transcriptional landscape were unraveled by comparing the global expression profiles of all the expressed genes. RNA-Seq technique was used to study alternative gene expression in *M. smegmatis* WT, *ΔdisA*, and *Δpde* strains (42). Since c-di-AMP is known to be produced at a basal level under normal physiological conditions (**Fig.S2**), cells were grown till the mid-log phase, harvested to isolate intact RNA, and then performed the RNA-seq analysis. Differential expression analysis was carried out using the DESeq2 package and genes (more than a total of 5 reads) with absolute log2 fold change≥1 with p value≤0.05 were considered to be significant. Out of a total of 6518 expressed genes for all strains, in *M. smegmatis* WT (Reference) vs *M. smegmatis ΔdisA* (Test) comparison, the total number of significantly altered genes was found to be 327. Similarly, *M. smegmatis* WT (Reference) vs *M. smegmatis Δpde* (Test) comparison revealed a total of 109 genes were differentially expressed (Up- and down-regulated **(Fig.5A)**. The expression profile of the differentially expressed genes across the samples was presented in volcano plots **(Fig.5B)**. The genes that showed significant differential expression (highlighted in volcano plot with statistical significance) were used for Gene Ontology (GO) and pathway enrichment analysis. Next, we performed enrichment analysis of genes in 3 following categories: Biological process (BP), Molecular function (MF), Cellular component (CC), and KEGG Pathway using ClusterProfiler R software **(Fig.5C)**. Gene Ontology (GO) and pathway terms, with multiple test p-value ≤ 0.05 were considered significant in our analysis and illustrated in the Bubble plot **(Fig.5D)** where the size of the bubble is proportionate to the number of genes involved, different colors highlighted different categories of functional classifications. The orange baseline indicated the p-value threshold (Benjamini-Hochberg p-value 0.05). GO enrichment results revealed that in the case of *M. smegmatis ΔdisA* strain several ribosomal proteins and translational machinery genes were significantly downregulated, later we confirmed that the *ΔdisA* strain was moderately sensitive to several antibiotics targeting bacterial translation (data not shown). Apart from that, the Arginine biosynthesis pathway, FAD-binding, and some transcription factor genes were differentially expressed in the null c-di-AMP background (**Table 1**), which needs further investigation. In the case of *M. smegmatis Δpde*, several genes for integral membrane protein and inner membrane proteins are differentially expressed compared to WT possibly due to high c-di-AMP levels inside the cells. In addition to that, genes involved in extracellular pathways and cell wall synthesis were found to be significantly altered with a huge fold change in expression (**Table 1**). A large number of genes representing structural components of the cell wall and membrane porins could justify the contrasting surface properties of the *M. smegmatis Δpde* strain with *M. smegmatis* WT. Next, we shortlisted some candidate genes from importantly regulated pathways with significantly varied expression levels and correlated with the observed phenotype. The top two enriched KEGG pathways in *M. smegmatis ΔdisA* strain with significant genes were illustrated as representative images, where up and down-regulated genes were shaded red and green respectively **(Fig.S8)**.

**Table 1.** List of genes in different categories which were significantly downregulated in **(A)** *M.smegmatis ΔdisA* and **(B)** *M.smegmatis Δpde*.

| Sl. No. | Component/ Pathway /Function | Genes | Gene function |
| --- | --- | --- | --- |
| 1 | Ribosome structure and function | *rpsR2* | 30S ribosomal protein S18 |
| 2 |  | *rpmE* | 50S ribosomal protein L31 |
| 3 |  | *rpmH* | 50S ribosomal protein L34 |
| 4 |  | *rpsT* | 30S ribosomal protein S20 |
| 5 |  | *rplL* | 50S ribosomal protein L7/L12 |
| 6 |  | *rplI* | 50S ribosomal protein L9 |
| 7 |  | *rpmF* | 50S ribosomal protein L32 |
| 8 |  | *rpsP* | 30S ribosomal protein S16 |
| 9 |  | *rpmG1* | 50S ribosomal protein L33 |
| 10 |  | *pth* | Peptidyl-tRNA hydrolase |
| 11 | Response to stress | *MSMEG_5733* | Universal stress protein |
| 12 |  | *MSMEG_5245* | Universal stress protein |
| 13 |  | *MSMEG_4207* | Universal stress protein |
| 14 |  | *MSMEG_3950* | Universal stress protein |
| 15 | Arginine biosynthetic process | *argF* | Ornithine carbamoyltransferase |
| 16 |  | *argB* | Acetylglutamate kinase |
| 17 |  | *argD* | Acetylornithine aminotransferase |

B
| Sl. No. | Component / Pathway / Function | Genes | Gene function |
| --- | --- | --- | --- |
| 1 | Extracellular region and cell wall | *ripA* | Peptidoglycan endopeptidase RipA |
| 2 |  | *mspA* | Porin MspA |
| 3 |  | *mspB* | Porin MspB |
| 4 |  | *hup* | DNA-binding protein HU homolog |
| 5 | Sulfate transmembrane -transporting ATPase activity | *cyst* | Cysteine synthase B |
| 6 |  | *MSMEG_4533* | Sulfate-binding protein |
| 7 |  | *cysW* | Sulfate transport system permease protein |
| 8 | Monosaccharid e-transporting ATPase activity | *MSMEG_6018* | Xylose transport system permease |
| 9 |  | *MSMEG_6804* | Sugar ABC transporter substrate-binding protein |
| 10 |  | *MSMEG_4656* | Sugar ABC transporter ATP-binding protein |

**Figure 5.**
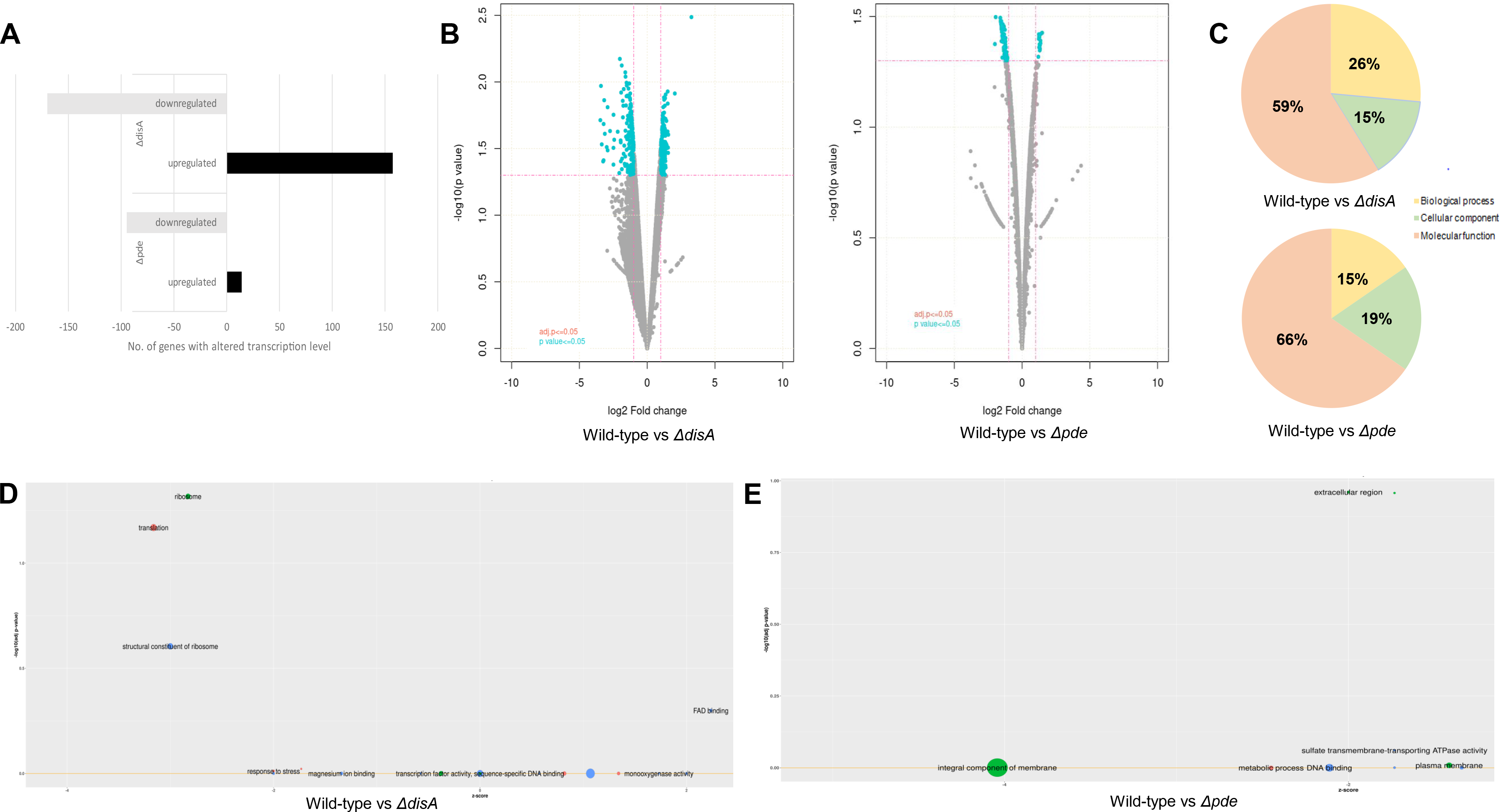
Transcriptome analysis by RNA-seq. **(A)** The number of differentially expressed genes in *M.smegmatis ΔdisA & M.smegmatis Δpde* **(B)** Volcano plot showing differential expression profile of genes between *M.smegmatis* WT and *M.smegmatis ΔdisA* (left) and *M. smegmatis* WT and *M.smegmatis Δpde* (right). The red line indicates log2 fold change≥1 and adjusted p value≤0.05. **(C)** The pie chart represents the pathway enrichment analysis of genes with only significant differential expression and is classified into 3 different categories: Biological process (BP), Molecular function (MF), Cellular component (CC) along with percentage abundance. Bubble plot depicts enriched Gene Ontology (GO) terms using significantly expressed genes in *M.smegmatis ΔdisA* **(D)** & *M.smegmatis Δpde* strains **(E)**. The orange line represents the p-value threshold (Benjamini-Hochberg p-value 0.05). The size of the bubble is proportionate to the number of genes involved in the GO term.

## Discussion

In our study, we have sought to understand the role of c-di-AMP in *M. smegmatis* by comparing diverse phenotypes in strains with altered c-di-AMP levels. Unlike other second messengers in *M. smegmatis*, c-di-AMP synthesis and degradation have been carried out by two different enzymes resulting in strict regulation of steady-state homeostasis, which is critical for normal cell growth and maintaining important physiological functions (43). Since c-di-AMP is known to be dispensable in mycobacteria (14, 44), it gave us an ideal opportunity to individually delete the genes responsible for the c-di-AMP synthesis and degradation to check the precise relevance of the molecule during normal growth and under relevant stress conditions **(Fig.6)**. So far 5 major types of deadenylate cyclase (DAC) enzymes have been discovered, which are DacA, DisA, CdaACdaM, CdaS, and CdaZ (45). Though the DisA class of DAC enzymes share the common N-terminal DAC domain with other more phylogenetically abundant classes like CdaA, in addition, it possesses a C-terminal DNA binding helix-hairpin-helix (HhH1) domain for monitoring DNA damage (2). In *B. subtilis*, DisA has been shown to be involved in sporulation control (3), however, this specific function of DisA seemed not widespread as many spore formers bacteria do not contain the *disA* gene in the genome. Since *M. smegmatis* is not known to form endospores and DisA is the sole c-di-AMP synthase, it was important to reveal other molecular functions regulated by c-di-AMP and DisA and understand how c-di-AMP synthesis and DNA damage response functions of the DisA enzyme in *M. smegmatis* gets coordinated.

**Figure 6.**
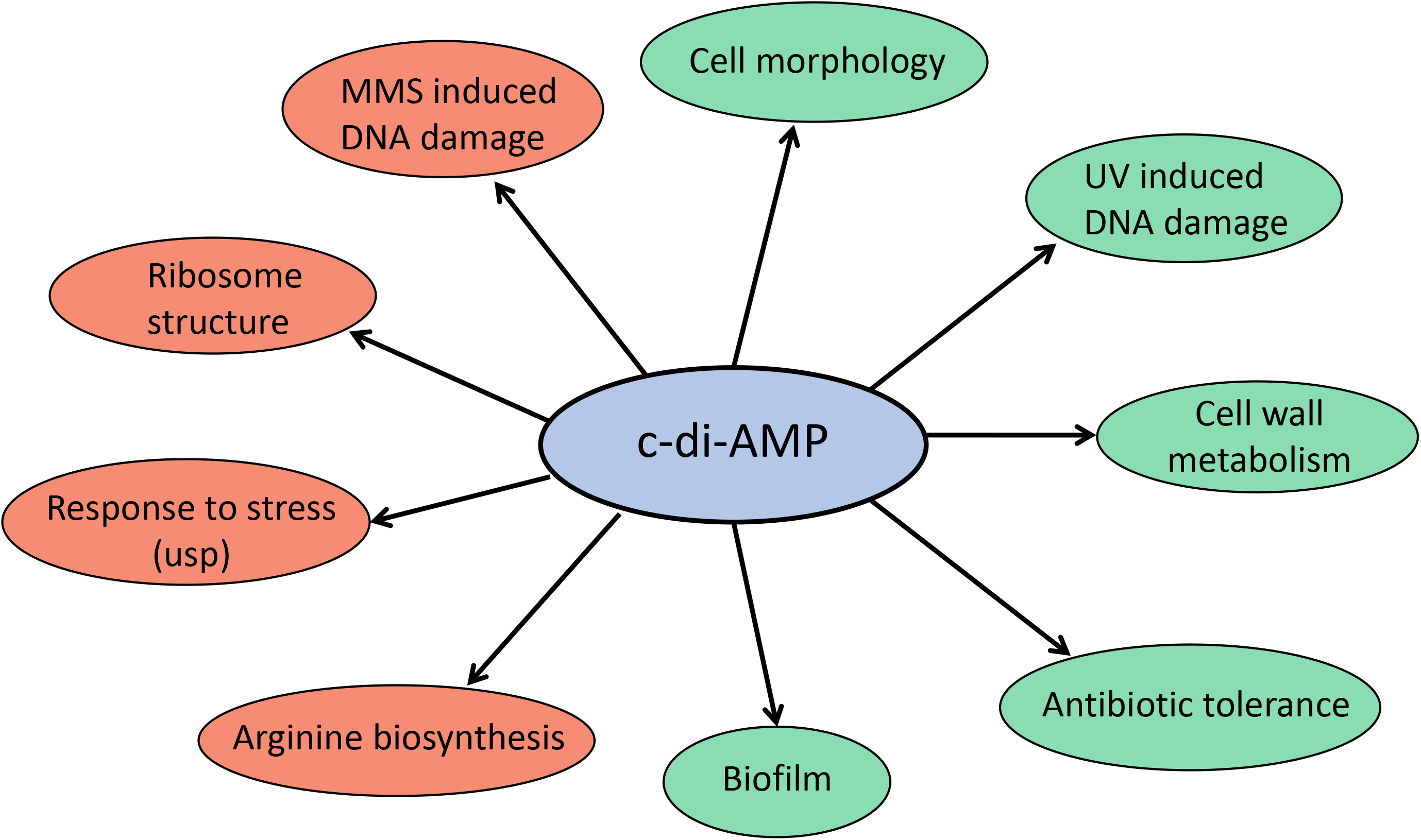
Schematic representation showing different cellular pathways regulated by c-di-AMP in *M.smegmatis*. The red and green bubbles depict c-di-AMP-null and overexpression phenotypes respectively.

We observed that c-di-AMP concentration affected several basic phenotypes like cell shape, cell length and cell width uniformity with visible but little differences in the strains’ growth profile. The colony appearances changed drastically with increased c-di-AMP concentration related to altered cell surface properties, which is further corroborated by different cellular aggregation profiles and surface hydrophobicity in *Δpde* strain. Further, we found that high c-di-AMP concentration does not favor the biofilm as well as air-liquid interphase pellicle formation. All these observations particularly for the *Δpde* strain could be attributed to major change in cell wall structure which was also pointed out by RNA-seq data.

Next, we checked *ΔdisA* and *Δpde* mutants’ survival patterns under different stresses and found that varying c-di-AMP concentration could play a definite role which is linked to underlying pathways yet to be discovered. The *Δpde* mutant was found to be very sensitive to UV irradiation compared to WT, possibly due to RecA enzyme’s functional deficiency (46) and the *ΔdisA* strain has better survival under identical conditions.

Another genotoxic stress MMS treatment made *ΔdisA* mutant extremely sensitive, which could indicate the relevance of DNA scanning properties of the enzyme DisA, not the c-di-AMP synthesis. SDS treatment revealed that both high and no c-di-AMP inside cells could be detrimental for the bacteria as possibly related to RNA-seq data where several cell wall and integral membrane protein synthesis genes/pathways seemed to be under the direct control of c-di-AMP. Next, by putting precise point mutations in different domains of DisA protein we could confirm that two domains of DisA have specific activity and could operate independently of each other.

Our Phenotypic Microarray analysis revealed how c-di-AMP concentration could play a major role in drug sensitivity by altering AUC values at specific antibiotic concentrations. In most of the cases, *Δpde* mutant showed hypersensitivity to different antibiotic classes with the diverse mechanism of actions (MOA) further corroborated by the difference in MIC and zone of inhibition (ZOE) studies. *M. smegmatis Δpde* is sensitive against ciprofloxacin, rifampicin, erythromycin and vancomycin. Further studies are already underway to discover the underlying molecular mechanisms of drug sensitivity related to the c-di-AMP level. On all occasions, *M. smegmatis ΔdisA* showed marginal resistance or no difference compared to WT implying the importance of maintaining optimum c-di-AMP concentrations inside cells. Previous studies have shown the direct link between increased c-di-AMP concentration with beta-lactam resistance in different gram-positive bacteria (4–5, 47–48), although the mechanism was not demonstrated. This effect only against cell wall inhibitors could be justified by a specific class of DAC enzyme (other than DisA) and its operon structure, known to directly modulate cell wall biosynthesis and osmolyte transport. In our case we observed the opposite result that the *Δpde* strain is sensitive to vancomycin compared to WT; which seemed more due to the downregulation of several genes in cell wall and extracellular factors as evidenced by RNA-seq data. The precise information about the pathway and mechanisms remains to be discovered in the future. Other than that, in *Listeria monocytogenes*, deletion of c-di-AMP synthase enzymes are linked to beta-lactam sensitivity, which again had some relationship with the intracellular localization of the membrane protein (49,50). In our case, *ΔdisA* showed hardly any modulation in drug sensitivity against different classes of antibiotics implicating there was no similar mechanism involved in *M. smegmatis*. Rather, our data suggest that *ΔdisA* is sensitive to aminoglycosides and marginally resistant to fluoroquinolone by diverse mechanisms (data not shown). Finally, RNA-seq based transcriptome analysis revealed alternative gene expression profiles in null- and overexpressing c-di-AMP mutants. Comparative analysis revealed 3-fold more genes were differentially expressed in *ΔdisA* mutant compared to *Δpde* mutant. Our analysis pointed out c-di-AMP driven regulation of several ribosomal structural genes which needs further investigation. Apart from that, several other pathways were shown to be affected in c-di-AMP null strain including efflux pump regulation, arginine biosynthesis and FAD binding. In the case of *Δpde* mutant, several components of the cell wall and membrane biosynthesis were affected due to high c-di-AMP levels. Some of the key genes/pathways mainly related to antibiotic response, regulated by c-di-AMP which were first identified by phenotypic microarray, then correlated with RNA-seq analysis were further verified by us through other assays along with complementation constructs. Important mechanistic insights revealed that c-di-AMP acts as a crucial determinant for regulating drug susceptibility/ resistance phenotypes in *M. smegmatis* (unpublished data). All in all, our study has revealed many novel phenotypes related to c-di-AMP in mycobacteria. Our ongoing and future research will bring out more interesting findings and hopefully further translate this crucial knowledge to understand *M. tuberculosis* physiology.

## Acknowledgment

AG thanks the Ramalingaswami Re-entry fellowship and Department of Biotechnology (DBT), Government of India, for funding this work (BT/RLF/Re-entry/31/2017). VC, AP and MS acknowledge the Department of Biotechnology (DBT), Government of India for their fellowships.

We acknowledge Dr. Avisek Mahapa for his help in handling the Omnilog PM system, which was bought with a grant from the Department of Biotechnology (DBT), Government of India. We acknowledge Mr. Sudhansu Gautam for his help in sample analysis with HPLC.

We thank Dr. Sushma Krishnan, Indian Institute of Science and Dr. Preeti Bhardwaj, NCBS, Bangalore for performing TEM and SEM microscopy respectively.

We thank Prof. Dipankar Chatterji, MBU, IISc for valuable feedback on the work.

VC and AG contributed to the conception and design of the study. VC, AP, MS and AG performed the experiments. VC, AP, MS and AG participated in data analysis and interpretation. VC, AP and AG wrote the manuscript.

The authors declare no conflict of interest.

## Supplementary file details

Figure S1, S2, S3, S4, S5, S6, S7, S8. Table S1, S2, S3.

