## Supplementary data for "Elucidating the role of c-di-AMP in *Mycobacterium smegmatis*: phenotypic characterization and functional analysis"

### Supplementary materials

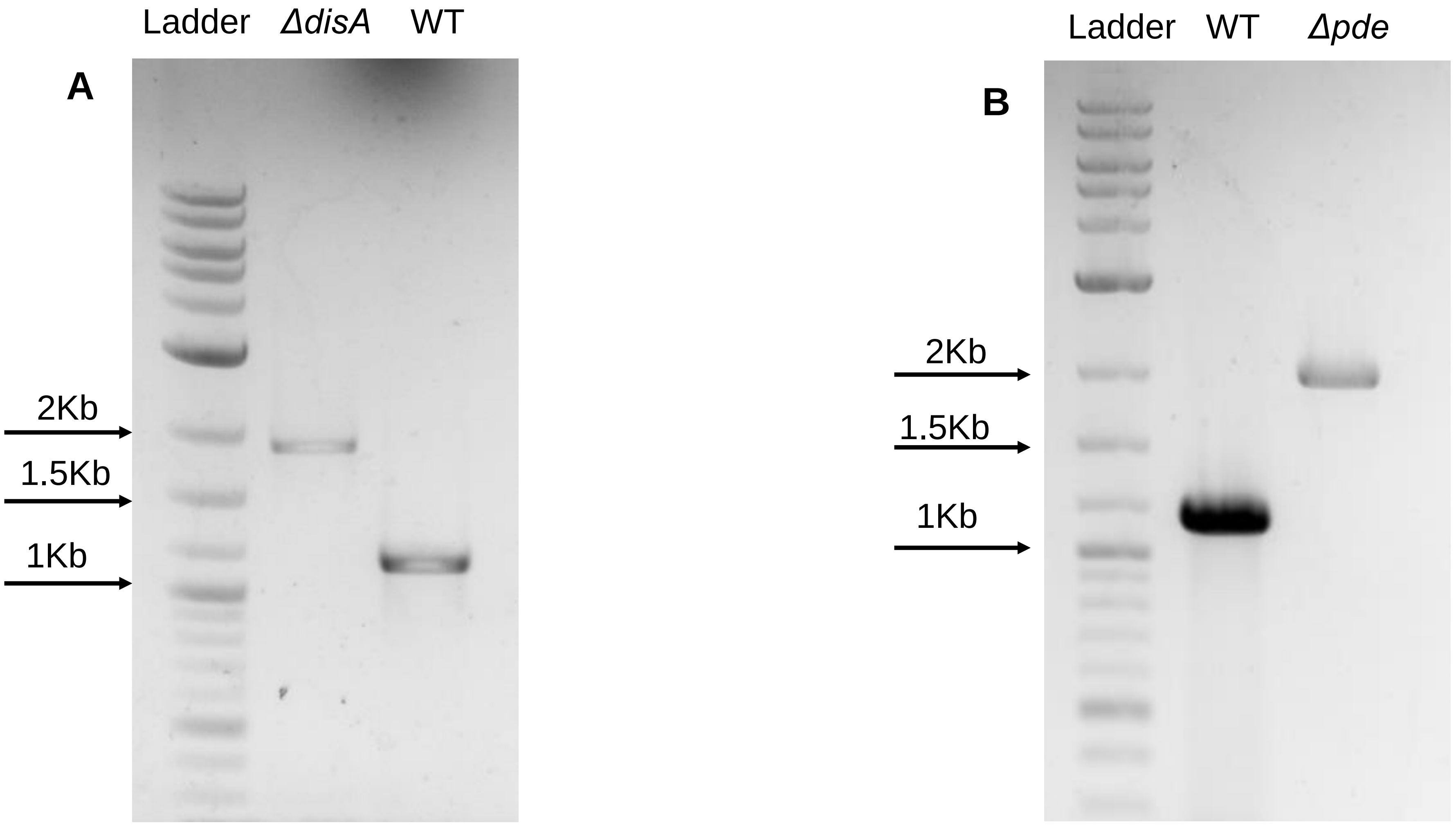

**Figure S1:** PCR confirmations of deletion of **(A)** *disA* and **(B)** *pde* genes in respective mutants

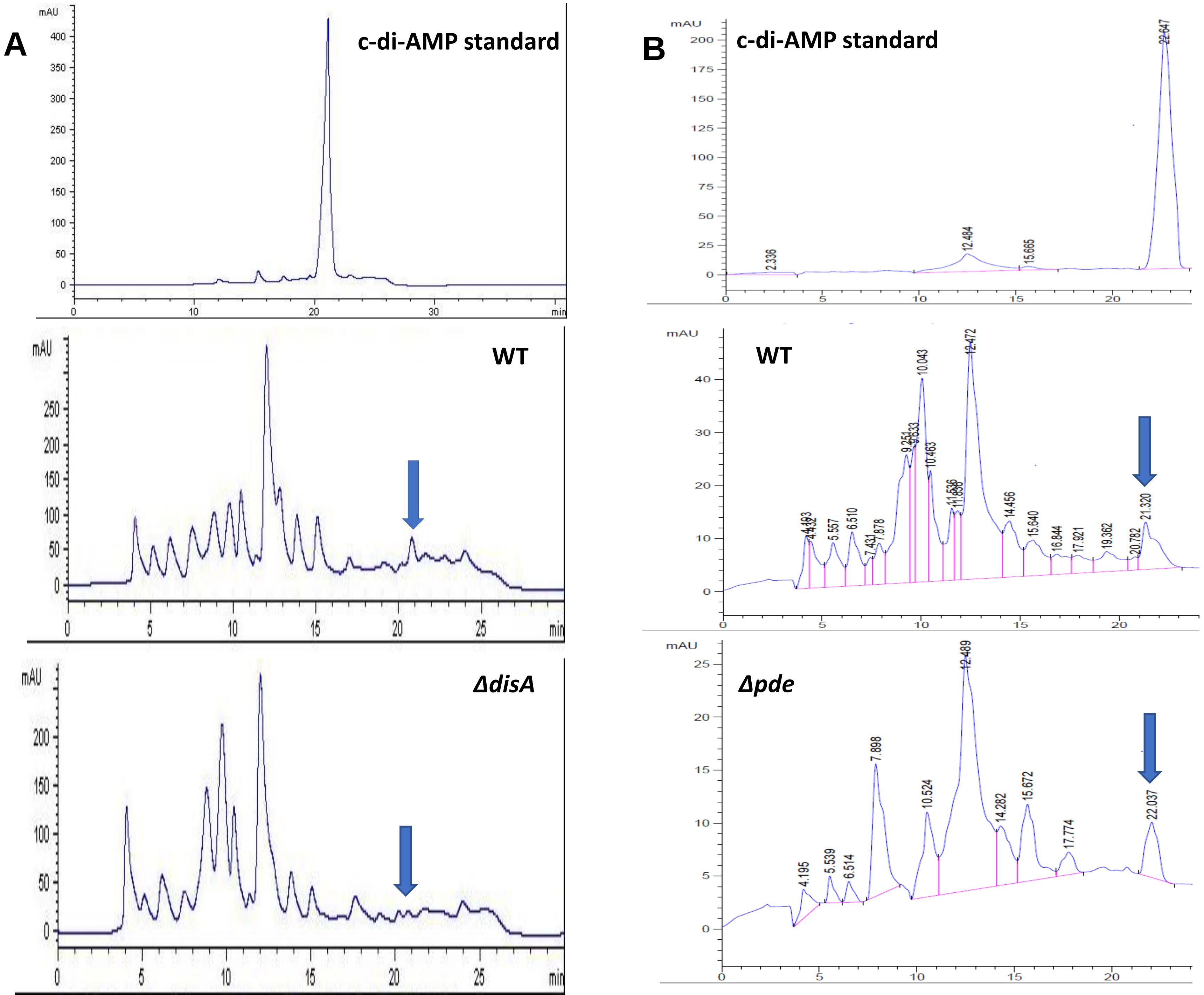

**Figure S2:** Detection of c-di-AMP from total nucleotide extract by reverse phase-HPLC. **(A)** c-di-AMP standard, WT and *ΔdisA* strain; **(B)** c-di-AMP standard, WT and *Δpde* strain Arrows indicate the position of the c-di-AMP peaks in different strains.

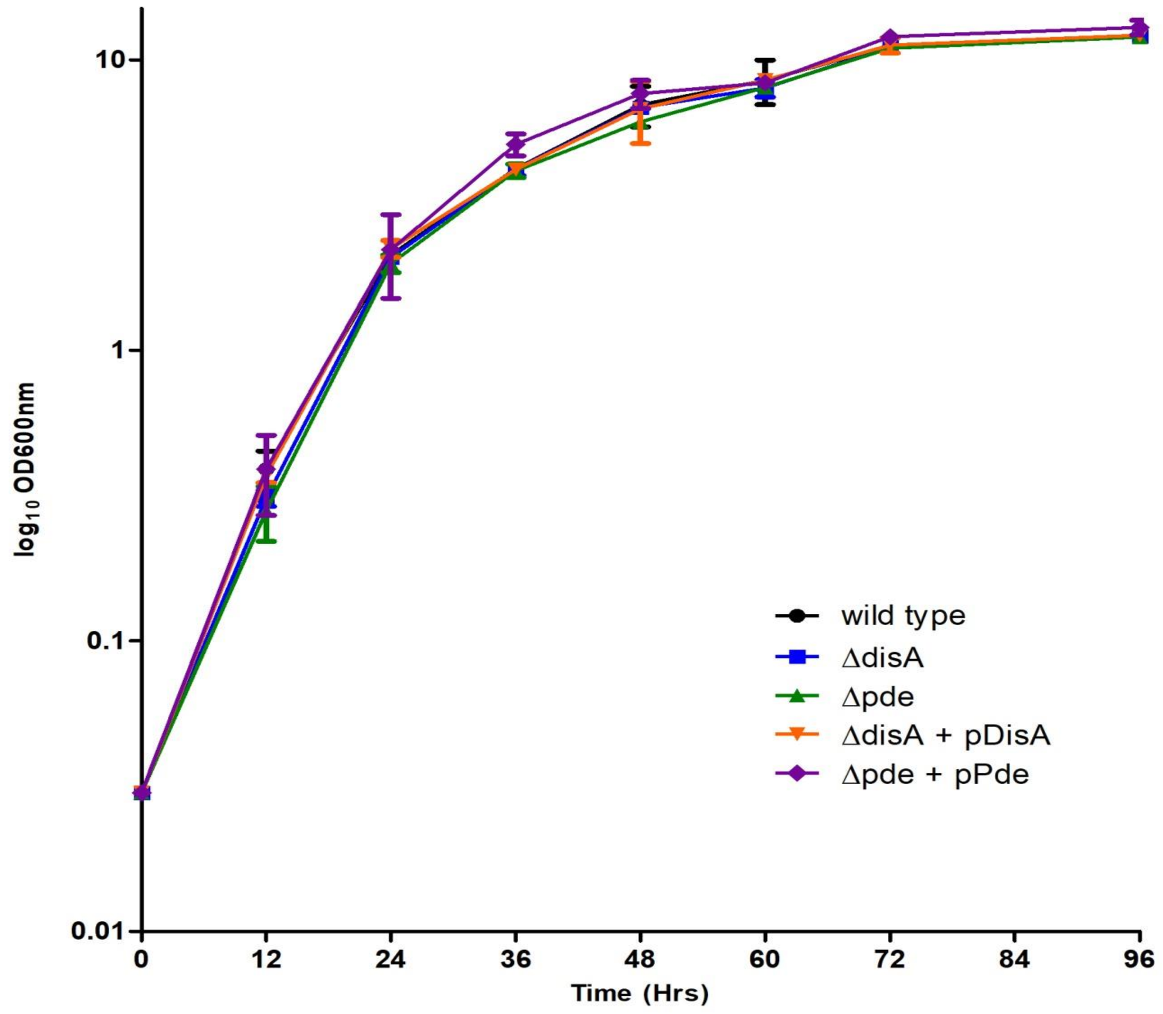

**Figure S3:** Representative growth curves of WT, deletion mutants and complementation strains in MB 7H9 containing 2% glucose and 0.05% Tween 80 with appropriate antibiotics. Growth profiles were monitored for 96 hours by measuring the optical density at 600nm using a spectrophotometer.

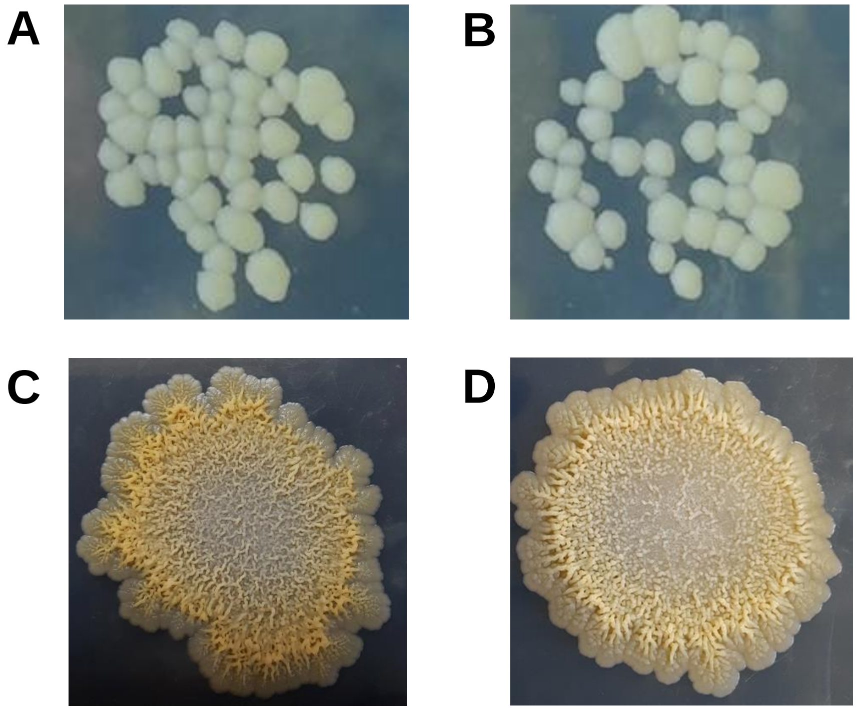

**Figure S4:** Colony morphology and colony architecture of *M.smegmatis ΔdisA* + pDisA strain **(A and C)** and *Δpde* + pPde strain **(B and D)**, similar to WT.

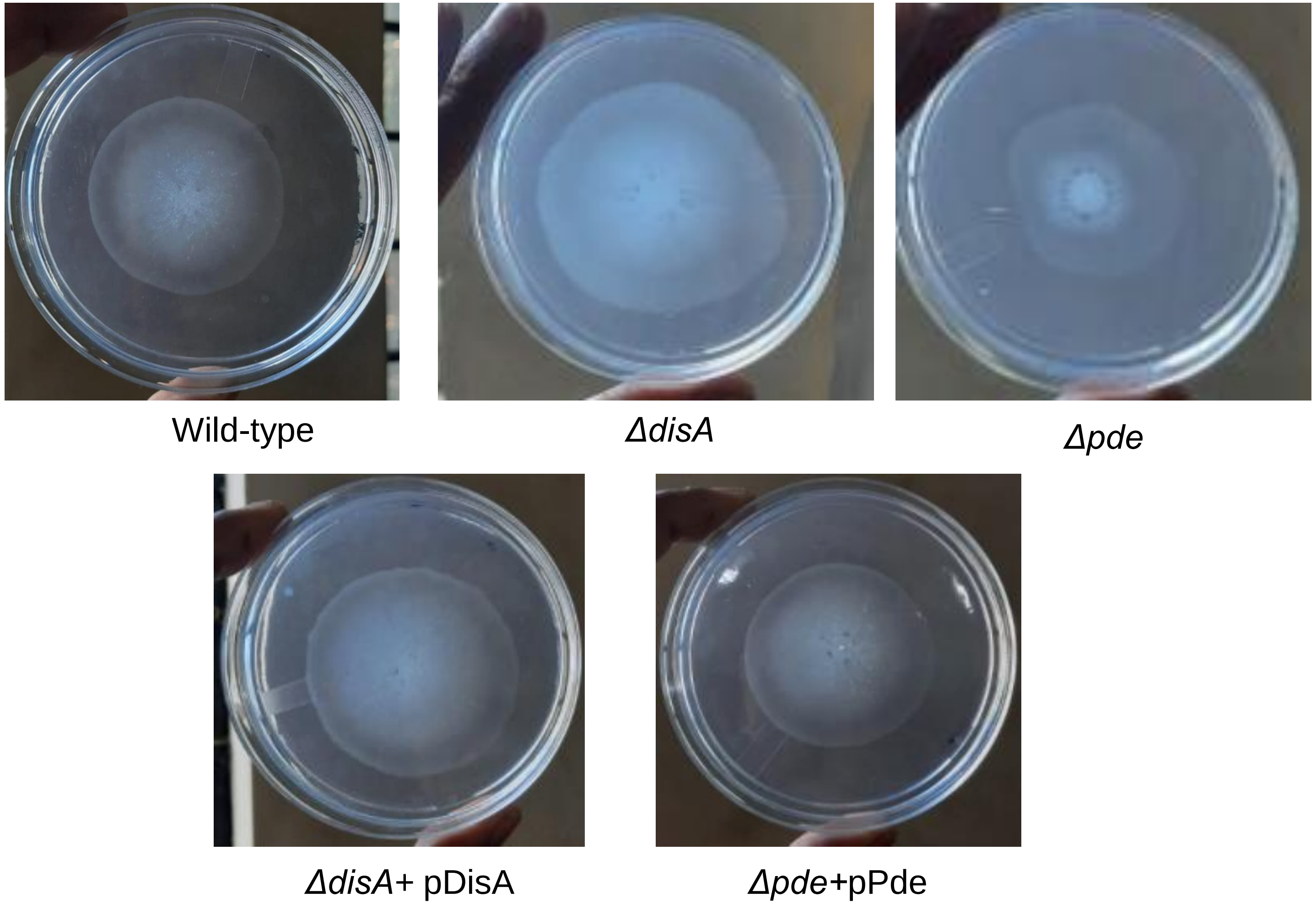

**Figure S5:** Different sliding motility pattern of different strains was observed after 3 days of incubation at 37 deg.

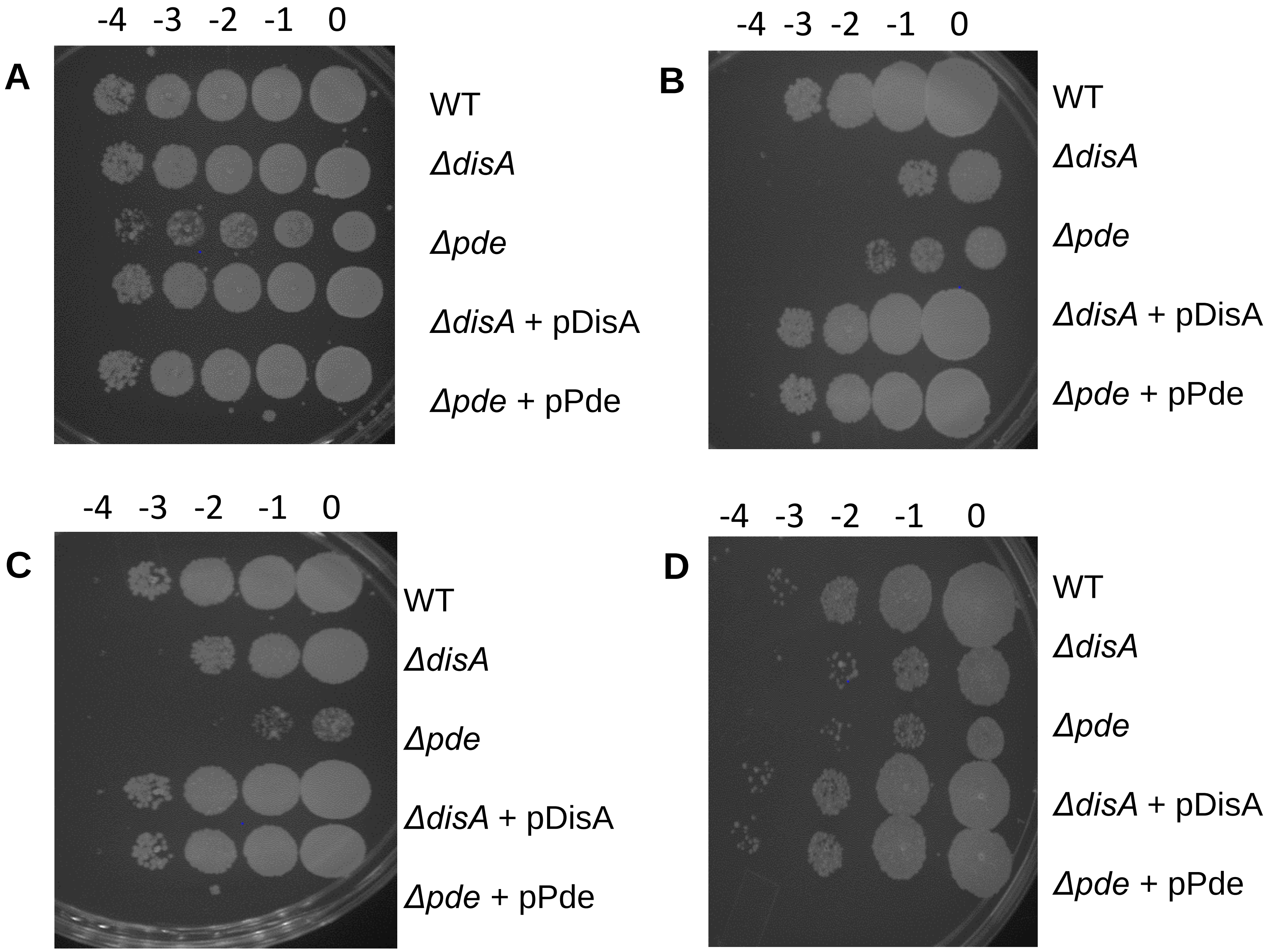

**Figure S6:** Representative spot plate pictures of **(A)**Untreated,**(B)** MMS, **(C)** UV and **(D)** SDS treated samples.

| PM plate wells | Antibiotic | Mode of action | Difference in AUC | |
|---|---|---|---|---|
| | | | WT/*ΔdisA* | WT/*Δpde* |
| Plate 20B | Ciprofloxacin | DNA replication | 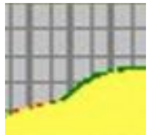 | 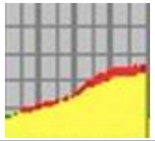 |
| Plate 12B | Rifampicin | Transcription | 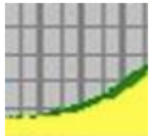 | 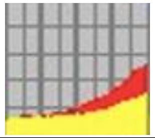 |
| Plate 11C | Erythromycin | Translation | 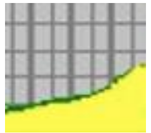 | 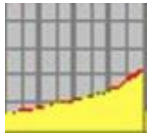 |
| Plate 12B | Vancomycin | Cell wall | 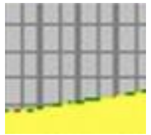 | 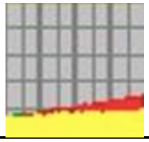 |

**Figure S7:** PM AUC curves highlighting differential growth pattern in presence of selective antibiotics

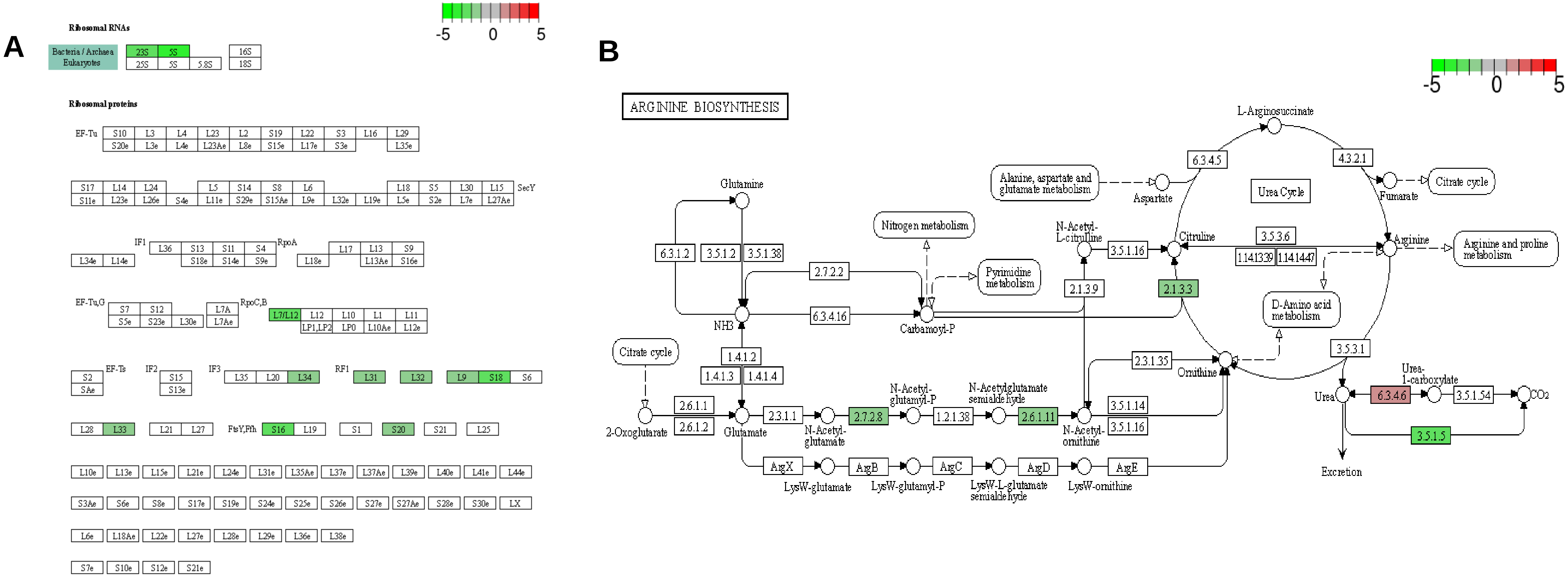

**Figure S8:** Representative images of top enriched KEGG pathways in *ΔdisA* strain highlighting the significantly up and down-regulated genes (shaded in red and green respectively): **(A)** Ribosome structure/function, **(B)** Arginine biosynthesis. The scale is set as log2fold change.

A

| Sl. No. | Antibiotics/Chemicals | Difference in AUC |
|---|---|---|
| **1** | Paromomycin | 33391.00 |
| **2** | Patulin | 31422.00 |
| **3** | Neomycin | 27646.00 |
| **4** | Chloroxylenol | 24845.00 |
| **5** | Iodonitro Tetrazolium Violet | 19668.00 |
| **6** | Aluminum Sulfate | 19013.00 |
| **7** | Ciprofloxacin | 18867.00 |
| **8** | Harmane | 16887.00 |
| **9** | Alexidine | 15994.00 |
| **10** | Blasticidin S | 13461.00 |
| **11** | L-Glutamic-g-Hydroxamate | 13143.00 |
| **12** | Methyltrioctylammonium Chloride | 11622.00 |
| **13** | Gentamicin | 11575.00 |
| **14** | Domiphen bromide | 11505.00 |
| **15** | Hydroxylamine | 11434.00 |
| **16** | Tinidazole | 11077.00 |
| **17** | Trimethoprim | 11006.00 |
| **18** | Sanguinarine | 10800.00 |
| **19** | Phosphomycin | 10560.00 |
| **20** | Chlorambucil | 9848.00 |
| **21** | Ampicillin | -29077.00 |
| **22** | Antimony (III) chloride | -25706.00 |
| **23** | Oxolinic acid | -25229.00 |
| **24** | Puromycin | -25148.00 |
| **25** | Cefsulodin | -24527.00 |
| **26** | Cefuroxime | -24178.00 |
| **27** | Ornidazole | -21532.00 |
| **28** | Azlocillin | -19839.00 |
| **29** | Thiosalicylate | -18374.00 |
| **30** | 5,7-Dichloro-8-hydroxy-quinaldine | -17313.00 |
| **31** | Poly-L-lysine | -16237.00 |
| **32** | Cesium chloride | -16215.00 |
| **33** | Compound 48/80 | -16145.00 |
| **34** | 3, 4-Dimethoxybenzyl alcohol | -15122.00 |
| **35** | Fusaric Acid | -14954.00 |
| **36** | Sodium periodate | -13258.00 |
| **37** | 5-Chloro-7-Iodo-8-Hydroxyquinoline | -13128.00 |
| **38** | Hydroxyurea | -12859.00 |
| **39** | Thallium (I) acetate | -12817.00 |
| **40** | 6-Mercaptopurine | -12594.00 |

**B**

| Sl. No. | Antibiotics/Chemicals | Difference in AUC |
|---|---|---|
| 1 | Paromomycin | 38525.00 |
| 2 | Neomycin | 27406.00 |
| 3 | Chloroxylenol | 20437.00 |
| 4 | Iodonitro Tetrazolium Violet | 12772.00 |
| 5 | Cefotaxime | 11623.00 |
| 6 | Demeclocycline | 11239.00 |
| 7 | Hydroxylamine | 9855.00 |
| 8 | Lincomycin | 9805.00 |
| 9 | Plumbagin | 9639.00 |
| 10 | Procaine | 8960.00 |
| 11 | Lidocaine | 8117.00 |
| 12 | Nickel chloride | 7820.00 |
| 13 | Penicillin G | 7687.00 |
| 14 | Blasticidin S | 7394.00 |
| 15 | Minocycline | 7330.00 |
| 16 | Zinc chloride | 7324.00 |
| 17 | Sulfanilamide | 7186.00 |
| 18 | Sodium azide | 6977.00 |
| 19 | Sanguinarine | 6740.00 |
| 20 | Alexidine | 6561.00 |
| 21 | Methyltrioctylammonium Chloride | -29121.00 |
| 22 | Antimony (III) chloride | -25140.00 |
| 23 | Aminotriazole | -23890.00 |
| 24 | Oxolinic acid | -23139.00 |
| 25 | EGTA | -22157.00 |
| 26 | Oxytetracycline | -22101.00 |
| 27 | Aminotriazole | -21673.00 |
| 28 | 3,5- Diamino-1,2,4-triazole (Guanazole) | -21393.00 |
| 29 | Patulin | -21288.00 |
| 30 | 6-Mercaptopurine | -20276.00 |
| 31 | Ornidazole | -19993.00 |
| 32 | Oxolinic acid | -19675.00 |
| 33 | 4-Hydroxycoumarin | -19618.00 |
| 34 | Ruthenium red | -19555.00 |
| 35 | Iodoacetate | -19132.00 |
| 36 | D,L-Methionine Hydroxamate | -18143.00 |
| 37 | Hexaminecobalt (III) Chloride | -17451.00 |
| 38 | Cefsulodin | -17267.00 |
| 39 | D-Serine | -16592.00 |
| 40 | Azlocillin | -16432.00 |

**Table S1:** List showing Antibiotics/ Chemicals (top 40) with difference in AUC values in **(A)** WT/ *ΔdisA* and **(B)** WT/ *Δpde.* ‘+‘ and ‘-‘ sign indicates comparative resistance and sensitivity respectively.

| **Strains/ Plasmids** | **Description** | **Source/ Reference** |
|---|---|---|
| pPR27 | Suicide vector, GenR | Pelicic *et al*. (1997) |
| pPR27_us-kan-ds_disA | Suicidal vector construct to delete chromosomal copy of *disA*, GenR, KanR | This study |
| pPR27_us-kan-ds_pde | Suicidal vector construct to delete chromosomal copy of *pde*, GenR, KanR | This study |
| pMV261_Empty | Cloning vector with hsp60 promoter, KanR | Stover *et al*. (1991) |
| pMV361_Empty | Integrative vector used for complementation, HygR | Stover *et al.* (1991) |
| pMV361_*disA* | *disA* cloned in pMV361 vector, HygR | This study |
| pMV361_*pde* | *pde* cloned in pMV361 vector, HygR | This study |
| pET28a_*disA*(D84A) | *disA*(D84A) cloned in pET28a vector, KanR | Gautam *et al.* (2021) |
| pMV261_*disA*(D84A) | *disA*(D84A) cloned in pMV261 vector, KanR | This study |
| pMV261_*disA*(R353A, R356A) | *disA*(R353A, R356A) cloned in pMV261 vector, KanR | This study |
| pMV361_*disA*(D84A) | *disA*(D84A) cloned in pMV361 vector, HygR | This study |
| pMV361_*disA*(R353A, R356A) | *disA*(R353A, R356A) cloned in pMV361 vector, HygR | This study |
| *E. coli* DH5α | *E. coli* host strain for DNA cloning | Pharmacia Biotech |
| *M. smegmatis* Wild type(WT) | *M. smegmatis* mc2 155 | Snapper *et al*. (1990) |
| *M. smegmatis* Δ*disA* | *M. smegmatis* mc2 155 strain knock out for *disA*; KanR | This study |
| *M. smegmatis* Δ*pde* | M. smegmatis mc2 155 strain knock out for *pde*; KanR | This study |
| *M. smegmatis* Δ*disA*+pDisA | *M. smegmatis* Δ*disA* strain containing *disA* cloned in pMV361 vector; HygR,KanR | This study |
| *M. smegmatis* Δ*pde*+pPde | *M. smegmatis* Δ*pde* strain containing *pde* cloned in pMV361 vector; HygR.KanR | This study |
| *M. smegmatis* Δ*disA*+Pmv361 Empty | *M. smegmatis* Δ*disA* strain containing empty vector pMV361; HygR,KanR | This study |
| *M. smegmatis* Δ*disA*+pDisA(D84A) | *M. smegmatis* Δ*disA* strain containing *disA* (D84A) in pMV361 vector; HygR,KanR | This study |
| *M. smegmatis* Δ*disA*+pDisA(R353A, R356A) | *M. smegmatis* Δ*disA* strain containing *disA*(R353A, R356A) in pMV361 vector; HygR, KanR | This study |

**Table S2:** Plasmids and strains used in the study

| Name | Sequence (5' to 3') | Description |
|---|---|---|
| MsdisA_us_Fw | ATCGTCTAGATCGAAGGCGACCGGGCATCG | To construct suicidal vector |
| MsdisA_us_Rev | AGCTGAATTCGGGTCGGGGACCAGCTGCAC | To construct suicidal vector |
| MsdisA_ds_Fw | ATGCGAATTCCGAGGAGCTCGACTCGCTCA | To construct suicidal vector |
| MsdisA_ds_Rev | CGATGCGGCCGCCGGGGTTGTCGCCGCGAT | To construct suicidal vector |
| MsPDE_us_Fw | ATCGTCTAGAGAGATCCGCGCCATCTTCCG | To construct suicidal vector |
| MsPDE_us_Rev | AGCTGAATTCTGCTCGGTATGTCAACGGTG | To construct suicidal vector |
| MsPDE_ds_Fw | ATGCGAATTCCGACGGCCCGGTTGCTGCCC | To construct suicidal vector |
| MsPDE_ds_Rev | CGATGCGGCCGCCGCAGCAGCAGATCCCGTCC | To construct suicidal vector |
| pJAM_disA_Fw | CGTAAGTACTATGGCCGTGAAGTCCGGCGC | To check the Δ*disA* knockout |
| pJAM_disA_Rev | ATGCTCTAGAGGCCAGCCGGTCGGCGATCG | To check the Δ*disA* knockout |
| pMV361_pde_Fw | ATTACAGCTGCGGGGACGAACGCTGAGGATG | To check the Δ*pde* knockout |
| pMV261_pde_Rev | TCAGAAGCTTGCATCAGCCAAGGGCCCGTGC | To check the Δ*pde* knockout |
| disA(D84A)_Fw | ACTGGAATTCATCCTAGGAGGGCTGATG | To construct pMV361_*disA*(D84A) |
| disA(D84A)_Rev | ACTGCAGCTGATCCTAGGAGGGCTGATG | To construct pMV361_*disA*(D84A) |
| disA(R353A, R356A)_Fw | TGTGGGCAGCCCACATCGCCGAAGGGCTCTCACTG | To construct pMV261_*disA*(R353A, R356A) |
| disA(R353A, R356A)_Rev | CCTTCGGCGATGTGGGCTGCCCACATCGAACCGATGC | To construct pMV261_*disA*(R353A, R356A) |

**Table S3:** Primers used in the study
